# Natural alteration of HIV-1 co-receptor tropism by contemporaneous HIV-2 infection

**DOI:** 10.1101/410456

**Authors:** Joakim Esbjörnsson, Fredrik Månsson, Hans Norrgren, Sarah L. Rowland-Jones

## Abstract

In this study, we show that the pathogenic HIV-1 CXCR4-tropism is more common in HIV-1 single (79%) than in HIV-1 and HIV-2 dual-infected individuals (35%), suggesting that contemporaneous HIV-2 infection can affect HIV-1 co-receptor tropism in late-stage disease. Understanding the underlying mechanisms responsible for this natural alteration by HIV-2 could pave the way towards a deeper understanding of the AIDS pathogenesis.

---

More than 25 million individuals have died of HIV-related causes since the discovery of HIV in 1983. Despite tremendous efforts, there is no cure or effective vaccine against the virus. The natural course of an HIV infection is usually described by three stages. The acute infection is characterized by viremia, rapid decrease in CD4^+^ T-cell counts and flu-like symptoms. In the asymptomatic stage, viral load is generally low and the CD4^+^ T-cell decline moderate. Finally, in the AIDS stage, viral loads increase, the CD4^+^ T cell count continues to decrease and opportunistic diseases develop due to a dysfunctional immune system.

Two genetically related but distinct human lentiviruses, HIV-1 and HIV-2, have been described^1,2^. Whereas HIV-1 is pandemic, HIV-2 is mainly confined to West Africa. Both viruses share similar transmission routes, cellular targets and AIDS causatives. However, an HIV-2 infection is characterized by a much longer asymptomatic stage, lower plasma viral load, slower decline in CD4^+^T-cell counts, and lower mortality rate^3-6^.

HIV enters target cells via interactions with CD4 and a co-receptor, usually one of the chemokine receptors CCR5 or CXCR4. Whereas CCR5-using strains usually are present throughout the complete disease course, CXCR4-using strains generally emerge in late-stage disease, close to the AIDS onset, and is almost invariably associated with a subsequent increase in the rate of CD4^+^T-cell decline, accelerated disease progression, and a poor prognosis for survival^7^. The recent introduction of a new drug class interfering directly with the CCR5-use by HIV-1 has highlighted the clinical significance in understanding basic mechanisms involved in HIV-1 co-receptor evolution.

In West Africa, where both HIV-1 and HIV-2 is present, dual-infection with HIV-1 and HIV-2 has been reported with a prevalence of 0-3.2%^8,9^. Previously, we have shown that HIV-2 exerts a natural inhibition against HIV-1 disease progression, by comparing HIV-1 single with HIV-1 and HIV-2 dual-infected individuals^10-12^. Both survival-time and the time to develop AIDS were approximately two times longer in dual-infected individuals.

Here, we studied 28 HIV-1 single and 17 HIV-1 and HIV-2 dual-infected individuals to investigate differences in HIV-1 co-receptor tropism and genetic variation of HIV-1. All individuals were treatment-naïve and considered to be in late-stage disease, as defined by CD4+ T-cell count (≤200 cells/μl or ≤14%) or clinical AIDS (according to the CDC and WHO Disease Staging Systems). Clinical parameters were similar across the groups, with mean CD4^+^T-cell counts of 165 cells/mm^3^ (range 10-662) and CD4% of 10 (range 2-32) (**Table 1** and **Supplementary Table 1**). Blood plasma samples were collected from each individual and the tropism of HIV-1 was determined by a recombinant phenotypic assay as previously described^13^ (**Supplemetary Methods**).

**Table 1.** Clinical data of the single and dual-infected individuals.

|  | N<sup>a</sup> | Mean CD4<sup>+</sup><br>T-cell count<br>(range)<sup>b</sup> | Statistics<sup>c</sup> | Mean CD4%<br>(range) | Statistics<sup>c</sup> |
| --- | --- | --- | --- | --- | --- |
| Single-infected individuals | 28 | 153 (21-426) | 0.557 | 10 (2-20) | 0.347 |
| Dual-infected individuals | 17 | 181 (10-662) |  | 11 (2-32) |  |

<sup>a</sup>n = number of individuals. <sup>b</sup>Mean CD4<sup>+</sup> T-cell count measured in cell/mm<sup>3</sup>. <sup>c</sup>Differences
between the groups were tested statistically using the Student's T-test.

We found that 79% of the single-infected individuals had HIV-1 with CXCR4-tropism, whereas only 35% of the dual-infected individuals had HIV-1 of this phenotype (p=0.005, Fisher’s exact test) (**Supplementary Table 1**).

Several longitudinal studies have presented evidence of a positive correlation between HIV-1 diversity and the time after infection, during the asymptomatic stage^14-17^. Close to the onset of AIDS, the diversity generally stabilizes, or in some cases even decreases^15^. In a previous study, it was shown that the diversity (genetic variation of HIV-1 at comparable time-points) was significantly lower in dual than in single-infected individuals during the asymptomatic stage of infection^11^. Despite these differences, the diversity at the time of AIDS onset was similar between the groups, although the time to reach this level of diversity was different. This finding is in agreement with the “diversity threshold theory” that suggests that AIDS develops when the diversity exceeds a critical threshold^17,18^.

In light of these results, we set out to investigate differences in HIV-1 diversity between single and dual-infected individuals during late-stage disease. Phylogenetic analysis of 370 HIV-1 *env* V1-V3 clones (~940 bp) (mean 8.22 clones/individual) revealed that there where no significant difference in diversity during late-stage disease between single and dual-infected individuals (**Table 2**). The “diversity threshold theory” predicts that the total virus population grows unboundedly beyond this threshold^19^. Furthermore, once the threshold has been exceeded, selection would favour strains with high replication rate, even though slower growing strains also would expand their population sizes. This would most likely result in rapid evolution with a fluctuating and broad spectrum of HIV-1 diversity between different individuals, as seen in both single and dual-infected individuals (**Table 2**).

**Table 2.** Maximum likelihood estimates of HIV-1 sequence diversity for single and dual-infected individuals.

|  | n<sup>a</sup> | Mean Diversity<br>(range)<sup>b</sup> | Statistics<sup>c</sup> |
| --- | --- | --- | --- |
| Single-infected individuals | 28 | 19.8 (0.7-46.5) | 0.332 |
| Dual-infected individuals | 17 | 16.0 (0.7-45.5) |  |

<sup>a</sup>n = number of individuals. <sup>b</sup>Diversity is given in substitutions x 10<sup>-3</sup>/site. <sup>c</sup>Differences between the groups were tested statistically using the Student's T-test.

Although other hypotheses exist, several lines of evidence suggest that HIV-1 CXCR4-tropic viruses evolve from pre-existing CCR5-tropic viruses during the natural course of infection^20,21^. Mathematical modelling of evolution in HIV-1 co-receptor tropism have shown that the development of CXCR4-using strains is favoured by a weak immune system and depends on the level of specific antiviral responses against these strains^22^. It is possible that HIV-2 could alter the expression of cellular factors *in trans*, and thereby affect the HIV-1 co-receptor tropism in dual-infected individuals. *In vitro* studies have shown that HIV-2 infection generates higher levels of β-chemokines (the natural ligands of CCR5) in peripheral blood mononuclear cells than HIV-1 infection, and that this can inhibit HIV-1 infection and replication^23,24^. It was hypothesised that such up-regulation could favor HIV-1 to switch from CCR5 to CXCR4-use in dual-infected individuals at a higher rate than in single-infected individuals. In the current, we found the opposite, suggesting that the potential *in vivo* effect of increased β-chemokine levels does not result in higher levels of HIV-1 CXCR4-tropism in dual-infected individuals.

In summary, we show a lower prevalence of HIV-1 CXCR4-using tropism in dual than in single-infected individuals. Our results suggest that the “diversity threshold theory” is a plausible model for HIV-1 evolution also in late-stage disease. Alterations in co-receptor tropism could be an important correlate of the natural inhibition against HIV-1 disease progression by contemporaneous HIV-2 infection. Further investigations of the interplay between HIV-1 and HIV-2 could reveal new and critical mechanisms towards a deeper understanding of AIDS pathogenesis.

## Acknowledgement

We thank Babetida N’Buna, Aquilina Sambu, Eusebio Ieme, Isabel da Costa, Jacqueline, Ana Monteiro Watche, Cidia Camara, Braima Dabo (deceased), Carla Pereira, Julieta Pinto Delgado, Leonvengilda Fernandes Mendes, Ana Monteiro, Ansu Biai, Helen Linder, Wilma Martinez-Arias, Elzbieta Vincic, Patrik Medstrand and SCIBLU Genomics for their contributions to this work. We also thank the Swedish Research Council, the Swedish International Development Cooperation Agency/Department for Research Cooperation (SIDA/SAREC), the Swedish Society for Medical Research, the Crafoord Foundation, Lund, Sweden, the Royal Physiographic Society, Lund, Sweden, The Lars Hierta Memorial Foundation, Stockholm, Sweden, and Konsul Thure Carlsson Fund for Medical Research, Lund, Sweden for generous funding of this work.

## Supplementary Methods

### Sample set

The samples used in this study were selected from a sample set of 52 plasma samples from 52 HIV-1 infected individuals and was selected from a cohort of police officers from Guinea-Bissau, West Africa, based on sample availability and disease status. The cohort has been described in detail elsewhere^1-3^. In addition, five plasma samples from individuals with a recorded HIV-1 and HIV-2 dual-infection were included from a case-control cohort from Bissau, Guinea-Bissau^4^. Forty-seven of the samples were successfully amplified (29 from single-infected individuals and 18 from dual-infected individuals). Only individuals with subtype A-like HIV-1 strains could be analyzed in the recombinant phenotypic assay as described^5^. Two individuals were infected with HIV-1 of subtype C and CRF06_cpx, respectively, and were not subjected to further analyses (**Supplementary Table 1**). All of the individuals were treatment naïve and classified to be in late-stage disease, as defined by CD4+ T-cell count (≤200 cells/μl or ≤14%) or clinical AIDS (CDC: C or WHO: 4)^6,7^. In cases where more than one sample from late-stage disease was available, the last sample was chosen. Individuals diagnosed with tuberculosis and clinically categorized as CDC: C, but without other AIDS-defining symptoms were not included in the study. For patient samples DL2713H, DL2846I and DL3018H, there were no recorded CD4+ T-cell counts. These samples were included in the study based on previous observations of CD4+ T-cell counts of the same patient, according to the described criterions. Details of the plasma samples from Guinea-Bissau can be found in **Supplementary Table 1**.

### Amplification and sequencing

Viral RNA was extracted and purified from blood plasma samples, using RNeasy Lipid Tissue Mini Kit (Qiagen, Stockholm, Sweden) with minor modifications from the manufacturer’s instructions. Briefly, 200 μl of blood plasma was disrupted in 2000 μl Qiazol and 10 μg Carrier RNA (Qiagen). The aqueous phase was loaded onto a spin column by multiple loading steps. RNA was eluted in 40 μl of RNase-free water and treated with DNase I (Fermentas, Helsingborg, Sweden). Viral RNA was reverse transcribed using gene-specific primers, and the *env* V1-V3 region amplified using a nested PCR approach (The SuperScript™ III One-Step RT-PCR System with Platinum^®^ *Taq* DNA Polymerase and Platinum^®^ *Taq* DNA Polymerase High Fidelity, Invitrogen, Copenhagen, Denmark) according to the manufacturer’s instructions using primers JE12F (5’-AAAGAGCAGAAGATAGTGGCAATGA-3’) and V3A_R2 (5’-TTACAATAGAAAAATTCTCCTCYACA-3’) for one-step RT-PCR and E20A_F (5’-GGGCTACACATGCCTGTGTACCYACAG-3’) and JA169 for nested PCR^8^. The amplified V1-V3 region of approximately 940 base pairs (nucleotides 6430 to 7374 in HXB2; GenBank accession number K03455) was cloned using the InsTAclone cloning system (Fermentas) and TOP10 cells (Invitrogen). Twelve colonies were routinely picked from each sample and the cloned fragments were amplified with Platinum^®^ *Taq* DNA Polymerase High Fidelity (Invitrogen) using conventional M13 primers (−20 and -24). Individual clones were purified and sequenced using BigDye Terminator v1.1 Cycle Sequencing Kit (Applied Biosystems, Stockholm, Sweden) according to the manufacturer’s instructions using primers E20A_F and JA169^8^.

### Phylogenetic analysis

Sequences were assembled, and contigs were analyzed with CodonCode Aligner version 1.5.2 (CodonCode Corporation, Dedham, USA). Only sequences with open reading frames were subjected for further analysis. Subtype determination and removal of putative recombinants were performed as previously described. A final dataset of 370 HIV-1 *env* V1-V3 sequences (mean 8.22 clones/individual) were aligned using PRANK_+F_ with a neighbor joining tree, constructed in MEGA4, as guide tree^9,10^. The PRANK_+F_ algorithm aligns sequences using phylogenetic information and has been shown to align sequences in an evolutionarily sound way. The alignment was manually edited and codon-stripped, resulting in a final sequence length of 654 nucleotides. Maximum-likelihood (ML) phylogenies, based on 1000 bootstrap alignments, were constructed in Garli 0.951 (www.bio.utexas.edu/faculty/antisense/garli/Garli.html) ^11^. This method efficiently maximizes the tree loge likelihood by using a genetic algorithm implementing the nearest neighbor interchange (NNI) and the subtree pruning regrafting (SPR) algorithms on a random starting tree to simultaneously find and optimize the topology and branch lengths^11,12^. The diversity was calculated by averaging pairwise tree distances between patient-specific sequences. This was done in all of the generated trees, yielding 1,000 estimates for each individual. The median value for each individual was used in the analysis.

### Determination of coreceptor tropism

Coreceptor tropism was determined according to a previously described protocol^5^. Briefly, human kidney embryonic 293T cells and human glioma U87.CD4 cells, stably expressing CD4 and one of the chemokine receptors (CCR5 or CXCR4) were employed as indicator cells^13,14^. Chimeric viruses with patient-specific V1-V3 regions were generated based on the protocol from the Tropism Recombinant Test (TRT) with minor modifications^15,16^. Amplified V1-V3 fragments from each plasma sample and 43XCΔV, a NheI-linearized vector containing a full-length pNL4-3 genome with the V1-V3 region deleted, were transfected into 293T cells using the calcium phosphate precipitation method. Chimeric viruses were harvested and stored at -80°C. For infection, chimeric viruses were added in duplicate wells containing semi-confluent U87.CD4 cells. Cultures were analyzed at day one, seven and nine for p24 antigen production by ELISA (Biomérieux, Boxtel, The Netherlands).

### Statistics

All statistical analysis was performed using PASW Statistics 18, Release 18.0.0 (Polar Engineering and Consulting).

### Ethics

The study was approved by the Research Ethics Committee at the Karolinska Institute, Stockholm, and the Ministry of Health in Guinea-Bissau.

---

**Supplementary Table 1.** – Clinical parameters, HIV-1 subtype and tropism of the 47 analyzed study subjects.

| Patient No. | Status<sup>1</sup> | Sex<sup>2</sup> | CD4%<sup>3</sup> | CD4tot<sup>4</sup> | CDC<sup>5</sup> | WHO<sup>6</sup> | Subtype | Tropism<sup>7</sup> | Sample year |
| --- | --- | --- | --- | --- | --- | --- | --- | --- | --- |
| DL1996H | S | M | 5 | 157 | B | 3 | CRF02_AG | R5X4 | 2000 |
| DL2089J | S | M | 9 | 59 | B | 3 | CRF02_AG | R5X4 | 2003 |
| DL2096F | S | M | 5 | 22 | C | 4 | C | N/A<sup>8</sup> | 2003 |
| DL2249I | S | M | 2 | 21 | C | 4 | CRF02_AG | R5X4 | 2004 |
| DL2339E | S | M | 12 | 178 | B | 3 | CRF02_AG | R5X4 | 2003 |
| DL2365K | S | M | 9 | 133 | B | 3 | A3 | R5 | 2006 |
| DL2391G | S | M | 5 | N/A<sup>8</sup> | B | 2 | CRF02_AG | R5 | 2000 |
| DL2401M | S | M | 11 | 141 | B | 3 | CRF02_AG | R5X4 | 2004 |
| DL2713H | S | M | N/A<sup>8</sup> | N/A<sup>8</sup> | B | 2 | CRF02_AG | R5X4 | 2007 |
| DL2846I | S | F | N/A<sup>8</sup> | N/A<sup>8</sup> | B | 3 | A3 | R5X4 | 2005 |
| DL2853E | S | M | 11 | 137 | A | 1 | CRF02_AG | R5 | 1998 |
| DL2920H | S | M | 11 | 126 | B | 3 | CRF02_AG | X4 | 2004 |
| DL3018H | S | M | N/A<sup>8</sup> | N/A<sup>8</sup> | B | 3 | A3/CRF_02_AG | R5X4 | 2006 |
| DL3037E | S | M | 3 | 74 | B | 3 | A3/CRF_02_AG | R5X4 | 2005 |
| DL3039G | S | F | 7 | 148 | A | 2 | A3/CRF_02_AG | R5X4 | 2006 |
| DL3071H | S | F | 20 | 123 | B | 3 | A3/CRF_02_AG | R5X4 | 2005 |
| DL3087E | S | M | 4 | 62 | B | 2 | CRF02_AG | R5X4 | 2001 |
| DL3098I | S | F | 14 | 426 | N/A<sup>8</sup> | N/A<sup>8</sup> | CRF02_AG | R5X4 | 2007 |
| DL3169F | S | M | 9 | 315 | B | 3 | CRF02_AG | R5X4 | 2004 |
| DL3170F | S | M | 8 | 65 | B | 2 | CRF02_AG | R5X4 | 2000 |
| DL3234J | S | M | 10 | 216 | A | 2 | A3/CRF_02_AG | R5X4 | 2006 |
| DL3312E | S | M | 2 | 36 | C | 4 | CRF02_AG | R5X4 | 1998 |
| DL3633G | S | F | 8 | 112 | C | 4 | CRF02_AG | R5X4 | 2003 |
| DL3721C | S | M | 11 | 257 | A | 1 | A3/CRF_02_AG | R5X4 | 1997 |
| DL3733G | S | M | 19 | 137 | B | 3 | CRF02_AG | R5X4 | 2004 |
| DL4248G | S | F | 13 | 159 | B | 3 | A3 | R5 | 2005 |
| DL4477D | S | M | 14 | 141 | B | 3 | CRF02_AG | R5 | 2001 |
| DL4525G | S | M | 13 | 372 | B | 3 | A3 | R5 | 2006 |
| DL4632E | S | F | 9 | 77 | B | 3 | CRF02_AG | R5X4 | 2003 |
| DL2164G | D | M | 14 | 97 | B | 3 | A3 | R5X4 | 2005 |
| DL2198K | D | M | 32 | 145 | C | 4 | CRF02_AG | R5 | 2003 |
| DL2470F | D | M | 14 | 172 | C | 4 | A3 | R5X4 | 2001 |
| DL2544H | D | F | 17 | 59 | A | 1 | CRF02_AG | R5X4 | 2004 |
| DL2747I | D | M | 14 | 194 | B | 3 | CRF02_AG | R5X4 | 2005 |
| DL2829F | D | M | 9 | 183 | B | 2 | CRF06_cpx | N/A<sup>8</sup> | 2006 |
| DL3004H | D | M | 14 | 109 | B | 3 | CRF02_AG | R5 | 2003 |
| DL3247F | D | M | 4 | 108 | N/A<sup>8</sup> | N/A<sup>8</sup> | CRF02_AG | R5X4 | 2007 |
| DL3895F | D | M | 13 | 218 | B | 3 | CRF02_AG | R5 | 2001 |
| DL4084I | D | F | 12 | 442 | B | 3 | A1 | R5X4 | 2007 |
| DL4957C | D | F | 4 | 65 | B | 3 | A3 | R5 | 2007 |
| DL5342B | D | M | 2 | 133 | N/A<sup>8</sup> | N/A<sup>8</sup> | A3/CRF_02_AG | R5 | 2007 |
| DL6324B | D | M | 4 | 123 | B | 3 | CRF02_AG | R5 | 2007 |
| DL11967A | D | F | 13 | 440 | N/A<sup>8</sup> | N/A<sup>8</sup> | A3/CRF_02_AG | R5 | 2005 |
| DL11968A | D | F | 7 | 69 | N/A<sup>8</sup> | N/A<sup>8</sup> | A3 | R5 | 2005 |
| DL11969A | D | F | 12 | 55 | N/A<sup>8</sup> | N/A<sup>8</sup> | A3 | R5 | 2005 |
| DL11970A | D | F | 7 | 10 | N/A<sup>8</sup> | N/A<sup>8</sup> | A3/CRF_02_AG | R5 | 2006 |
| DL11971A | D | F | 11 | 83 | N/A<sup>8</sup> | N/A<sup>8</sup> | A3 | R5 | 2006 |

<sup>1</sup>S=HIV-1 single-infected individual. D=HIV-1/HIV-2 dual-infected individual. <sup>2</sup>M=male, F=female <sup>3</sup>CD4+ T cell percentage among all T cells <sup>4</sup>CD4+ T cell count per microliter among all T cells <sup>5</sup>Clinical category of the patient, as defined by the CDC, at the sample time point <sup>6</sup>Clinical category of the patient, as defined by the WHO, at the sample time point <sup>7</sup>R5=CCR5-tropic. X4=CXCR4-tropic. R5X4=dual-tropic (CCR5 and CXCR4-using). <sup>8</sup>N/A = not analyzed

